# Size Effect on Structure and Stiffness of Viral DNA during Temperature Variation

**DOI:** 10.1101/2020.08.29.273755

**Authors:** Cheng-Yin Zhang, Neng-Hui Zhang

## Abstract

Structure and stiffness of nucleic acids are closely related to viral infectivity. The article is aimed to clarify a recent controversy on structure variation of viral DNA in temperature-dependent experiments. A multi-scale correlation between microscopic structure of viral DNA and its bulk stiffness is formulated by presenting a two-zone mesoscopic free energy model. Results reveal that the increasing temperature promotes DNA structure transformation from order to disorder, and leads to size effect on temperature-dependency of stiffness and structure of viral DNA.

## Major Section

Viruses can infect almost all cellular life forms and cause diseases [1,2]. Researches reveal that virus infectivity is affected by its biological and chemical function as well as mechanical factors. By experiments of advanced technology, such as optical tweezers, atomic force microscopy (AFM) nanoindentation and small angle X-ray scattering (SAXS), virus structures and their bulk mechanical properties in vivo or vitro can be measured [3–8]. Kol et al. found that human immunodeficiency virus (HIV) particles during maturation got softening due to lack of some proteins, and soft particles entered cells more efficiently than stiffer ones [9]. Wilts et al. showed that a pH increase could primarily lead to the stiffness softening for the capsid of cowpea chlorotic mottle virus (CCMV) [10]. Gelbart’s group revealed that multivalent cations like Co^3+^ or spermine^4+^ could change DNA structure in λ phage, which suppressed DNA ejection from capsid [7,11]. These experiments show that changes in biological and chemical conditions can lead to variations of structure and bulk mechanical property of virus.

In addition to solution ion conditions, temperature is another significant environmental factor affecting viral infectivity [2]. Appropriate temperature ranges not only prolong the survival time of viruses and increase their infection incidences [12,13], but also affect their bulk mechanical properties. By light scattering experiments, de Frutos et al. and Löf et al. revealed an exponential increase in phage ejection rate with the increasing temperature [14,15]. As ambient temperature rises, by AFM nanoindentation experiments, Evilevitch et al. found the stiffness softening for the DNAs of herpes simplex virus type 1 (HSV-1) and λ phage, especially at the optimum temperature 37 °C [16–18]; simultaneously by SAXS, they discovered that the scattering peak underwent a prominent decrease, which indicated a remarkable transition of DNA structure in virus or phage capsid [16–18]. Livolant et al. tried to repeat the similar SAXS experiments on another four bacteriophages such as T5, λ, T7, and Φ29, however did not observe any significant change of the scattering area in the temperature range of 20–40 □, so they suggested that the temperature effect on viral DNA structure should be reconsidered [19]. So far the viewpoints on temperature effect of viral DNA structure seem controversial.

There is a difficulty to bridge the environment-dependent correlations between microscale structure and bulk mechanical property of soft active matters, especially on the micro/nano scale. To solve this problem requires the development of rules that translate molecular-scale characteristics to measurable mechanical responses [20]. Traditionally bulk DNA solution is thought to be a typical lyotropic liquid crystal, which is not sensitive to temperature. Here based on experimental results of viral DNA structure and the relevant mesoscopic/macroscopic theories of DNA liquid-crystalline, we manage to present a multi-scale temperature-dependent correlation between the microstructure of viral DNA and its bulk stiffness. First, based on the cryo-electron microscopy (cryo-EM) photos of viruses and phages [21,22], we propose a two-zone structural model of the packaged DNA in viral capsid, i.e., the occupied space of DNA is divided into an ordered zone and a disordered zone. Second, we update a mesoscopic energy model for the packaged DNA with the above two-zone structure by considering the contributions from microscopic interaction between adjacent DNA chains [23,24] and DNA intrinsic elasticity [25,26], then determine the two-zone structural variables with the minimum energy principle. Finally, we derive a bulk stiffness formulation of viral DNA in the AFM indentation with the reciprocal work theorem. Numerical results show that the temperature-dependence of viral DNA structure and its bulk stiffness has size-effect, and this not only clarifies the seemingly controversy about viral DNA structure by SAXS experiments, but also elucidates the mechanism of viral DNA stiffness change with temperature variation.

First, a two-zone structure assumption of viral DNA is proposed. Based on the cryo-EM observation on viral DNA structure [21,22], the space of viral DNA is divided into two regions, i.e. an outer tight ordered zone and an inner loose disordered zone as shown in Fig. 1. The radius of the entire structure, *R*_t_, namely the outer radius of the ordered DNA zone, is equal to the inner radius of the capsid; the inner radius of the ordered zone, *R*_oin_, is equal to the outer radius of the disordered zone, *R*_dout_; *R*_din_ is the radius of the innermost layer in the disordered DNA zone. In addition, *L* is the total length of the packaged DNA chain, *L*_ord_ is the DNA length in the ordered zone, and *L*−*L*_ord_ is that in the disordered zone. Illuminated by the two-phase coexistence theory of DNA liquid-crystalline proposed by Podgornik et al. [27,28], here we use two averaged interchain spacings to statistically distinguish two crystalline phases, i.e., line hexatic (LH) phase in the outer ordered DNA zone and cholesteric (LH) phase in the inter disordered DNA zone. And DNA conformation in each zone still follows the classical assumption of the inverse spool structure [25,26].

**FIG. 1.**
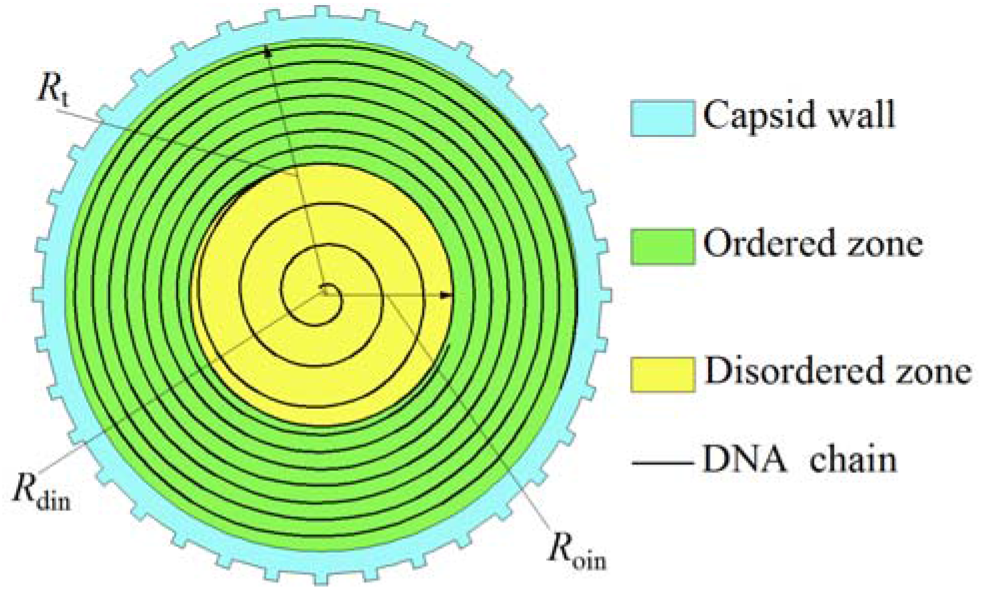
(color online) Cross section of viral DNA with two-zone structure.

Second, with the above assumption of two-zone structure, an updated mesoscopic energy model of viral DNA will be established. Considering the contributions from microscopic interaction between adjacent DNA chains [23,24] and the intrinsic elasticity in a bending DNA chain [25,26], the total free energy of the packaged DNA, *E*_total_, can be defined as the summation of the free energy in the ordered DNA zone and that in the disordered one, i.e.

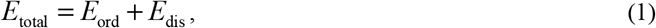

in which the subscript “ord” and “dis”, respectively, represent the ordered and disordered zones. The free energy in the ordered zone, *E*_ord_, and that in the disordered zone, *E*_dis_, respectively, are written as

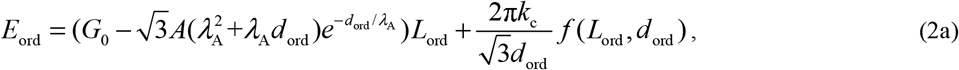

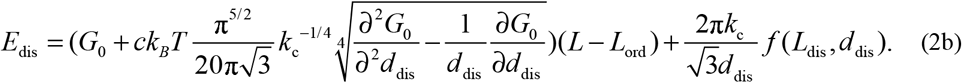

The first term on the right side of Eq. (2a) or (2b) represents the interactional energy among coarse-grained DNA cylinders [23,24], and the second term represents the elastic bending energy of a single DNA chain [25,26]. And *G*_0_(*d*) is the bare repulsive interactional energy per length in terms of interchain spacing *d*, including the contributions from the hydrated and electrostatic repulsion, written as

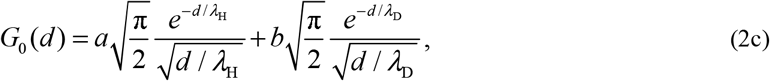

where the coefficients *a*, *b*, *c* and *A* are the hydrated, electrostatic, conformational entropic and attractive amplitude coefficient, respectively, *λ*_D_ is the Debye length, *λ*_H_ and *λ*_A_ are the hydrated length and attractive decay length, respectively. The values of *a*, *b*, *c*, *A*, *λ*_H_ and *λ*_A_ are obtained by fitting osmotic pressure experiment data [29], however Debye length *λ*_D_ has its own expression (see farther below Eq.(4)). *k*_c_ = *ξ*_p_*k*_B_*T* is the intrinsic bending stiffness of a single DNA chain, in which *ξ*_p_ is the persistence length of a DNA chain, *k*_B_ is the Boltzmann constant, *T* is the environmental temperature. Note that the difference in interactions between DNA strands in the two zones, by the similar method in Ref. [27,28], we update the interactional energy formulation of Parsegian et al. [23,24]. In the ordered zone, the conformational entropy are small and can be omitted due to the tight ordered arrangement of DNA strands, and the attraction between adjacent DNA strands due to van der Waals effect etc. should be considered. This kind of attraction, emerged with a negative term in the first term of Eq. (2a), can diminish the repulsion part.

Whereas in the disordered zone, the DNA arrangement is so loose and chaotic that the conformational entropy should be considered, so the DNA energy in the disordered zone is still described by the classical formulation of Parsegian et al., i.e. the first term in Eq. (2b) [23,24].

In addition to interaction energy of DNA strands, the contribution from bending elasticity of a DNA chain, expressed as the second term in Eq. (2a) or (2b), is also indispensable for the total energy. And *f*(*L*,*d*), as a function describing DNA structure, is in terms of chain length *L* and spacing *d* [25,26]. Substituting Eqs. (2a)–(2c) into Eq. (1) yields the free energy of the packaged DNA expressed as a function *E*_total_(*d*_ord_,*d*_dis_,*L*_ord_) of three unknown structural parameters, i.e. DNA length in ordered zone *L*_ord_, interchain spacing in the ordered zone *d*_ord_, spacing in the disordered zone *d*_dis_, and other given parameters.

Although bulk DNA solution was usually considered as a lyotropic liquid crystal, which is not sensitive to temperature change [30], some physical parameters in the above energy model are really temperature-dependent. The previous experiments revealed that the DNA persistence length is strongly dependent on temperature [31], and the following empirical formula was given

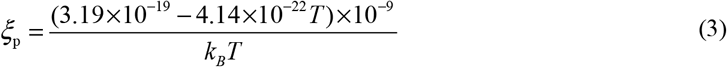

And the Debye length *λ*_D_, as an indispensable role in describing the electrostatic contribution, is also relevant to temperature, i.e

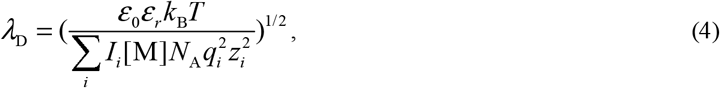

in which the solution dielectric constant *ε*_0_*ε*_r_ also depends on temperature [32]. These temperature-dependent physical properties in Eqs (3) and (4) actually are reflections of the temperature-dependence of DNA structure at the nanoscale from different profiles.

Next, the structural parameters, *d*_ord_, *d*_dis_, *L*_ord_, for the two-zone viral DNA can be determined by the minimum energy principle. As the above free energy function is too complicated to give an analytical relation, we turn to the numerical solution of structural parameters. To do this the energy *E*_total_(*d*_ord_,*d*_dis_,*L*_ord_) is first expanded into a second order Taylor’s expansion *E*_Taylor_(*d*_ord_, *d*_dis_,*L*_ord_), then the numerical solution can be obtained from the following functional stationary

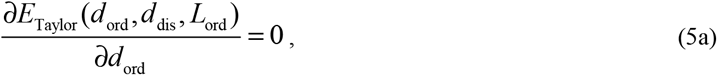

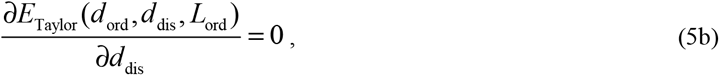

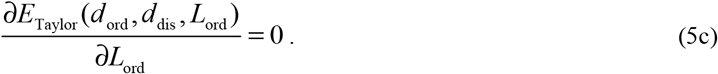

Note that the initial point choice for the Taylor’s expansion is important to numerical analysis, we first estimate the initial values of the three structural parameters in the Taylor’s expansion by referring to the data of DNA interchain distances of LH and CH phases in osmotic pressure experiment [27,28] and the actual structural observation by cryo-EM [21,22]. Then substituting the obtained numerical solutions of the three structural parameters into Eq. (2) yields the value of the free energy *E*_total_.

Third, after obtaining the relation between viral DNA structure and the corresponding free energy, we can calculate the bulk stiffness contributed by viral DNA in the AFM nano-indentation experiment. With the assumption of no energy dissipation during the indenting process and by the reciprocal work theorem, a relation between the bulk stiffness and the mesoscopic energy of viral DNA (see the details in Supplement 3) can be deduced. Out model reveals that the bulk stiffness of the entire particle is contributed by three parts: the capsid, the osmotic pressure and the DNA bending energy, i.e.

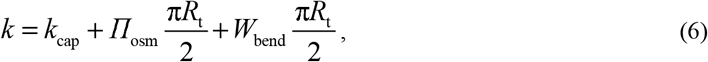

where

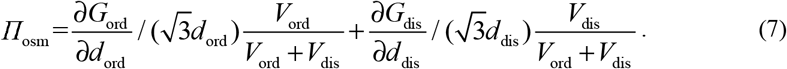

In Eqs. (6) and (7), *Π*_osm_ is the osmotic pressure induced by the packaged DNA, and *G*_ord_, *G*_dis_ are the interaction energy per unit length of DNA chain in the ordered zone or disordered zone, respectively. *W*_bend_ is the elastic energy density of DNA chain bending. Compared with the previous model of the pressure-enhanced viral stiffness [33], an extra contribution from DNA bending elasticity is included in the third term of Eq. (6).

In order to validate the two-zone structure assumption of viral DNA and its corresponding free energy model, λ phage is taken as a sample. The capsid geometry is treated as a spherical shell with an inner radius *R*_t_=29 nm, and the DNA length *L*=16500 nm [11]. Referring to the relevant experiments [34], we select 25mM Mg^2+^ and temperature *T*=298 K as solution condition [29]. By the present two-zone model, i.e. Eqs.(1) and (5), the structural parameters of the packaged DNA are predicted as: *d*_ord_=2.44 nm, *d*_dis_=2.72 nm, *L*=16130 nm, the number of DNA layers in the ordered zone is 4, and these predictions are consistent with the experiment observations by X-ray diffraction [35]. As for the packing energy consumed during the encapsulation of DNA into phage capsid, the prediction by the classical single-zone inverse spool model is 5.7×10^−5^ pJ [25], the present prediction is 6.79×10^−5^ pJ, while the detected value was 6.13×10^−5^ pJ when only 90% DNA completed its encapsulation [34]. Obviously, the present two-zone model is comparable to the classical single-zone model and has a more precise evaluation.

Next, HSV-1, λ phage and T7 phage are taken as examples to study temperature effect on the structure of viral DNA with the present two-zone model. The inner radius of HSV-1 capsid is 47 nm and its DNA length is 51500 nm [22]; the inner radius of T7 phage capsid is 22 nm and its DNA length is 14960 nm [21]; and 0.1 M Na^+^ is chosen as ionic conditions [29]. Fig. 2 shows the structural parameter variation of HSV-1 DNA with the temperature rising from 298 K to 314 K (the figures for the other two phages are shown in Fig. S1 of Supplement 1). As shown in Fig. 2 and Fig. S1, the DNA spacing in the disordered zone of T7 phage decreases slightly, whereas that in the ordered zone of T7 and both disordered zones of HSV-1 and λ phage enlarge with the increasing temperature. While the DNA length in the ordered zone is shortened as a whole with a non-monotonic local variation. These changes in structural parameters indicate that the rising temperature leads to an apparent rearrangement of DNA structure. Obviously the DNA length in ordered zone can be taken as a potential indicator to the transition of viral DNA structure from order to disorder due to the consistency of its changes with temperature variations for different viruses, whereas the single chain spacing is insufficient due to its diverse trends.

**FIG. 2.**
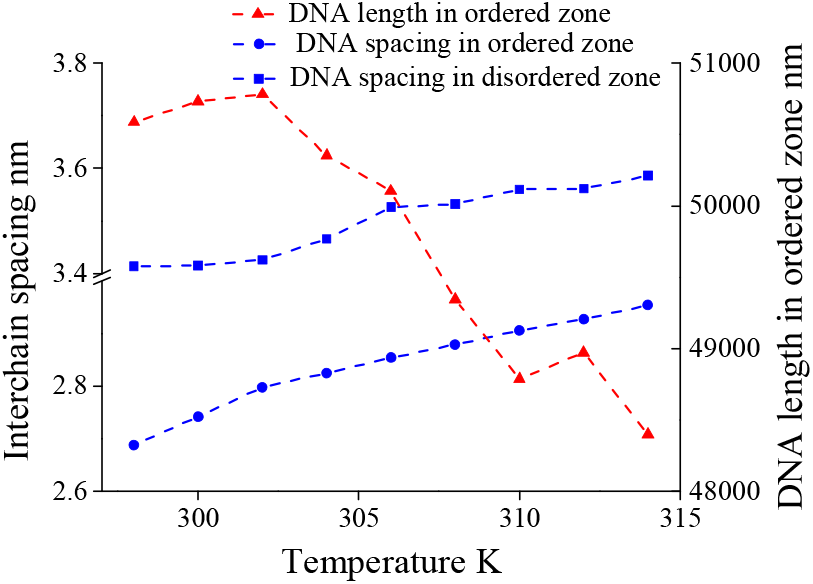
(color online) Structural parameter change with temperature for HSV-1 DNA under 0.1 M Na^+^ solution

In order to more directly characterize the structure change of viral DNA especially during phase-to-phase transition, we suggest the two-zone volume ratio as a potential indicator of the structural changes of the packaged DNA. Considering the surrounding prismatic space as shown in Fig S2 of Supplement 2, the effective DNA volume in each zone is written as

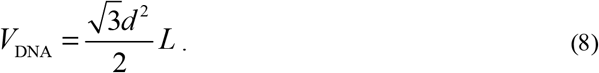

Fig. 3 shows that the two-zone volume ratios (*V*_ord_/*V*_dis_) all decline with the increasing temperature for three viral DNAs (HSV-1, λ phage and T7 phage). This indicates the relative enlargement of the disordered zone during the heating process, conforming to the common sense in thermal physics which temperature is related to the level of configurational disorder of a system [36]. Furthermore, the two-zone volume ratio of HSV-1 DNA with a larger capsid adopted in the experiment of Sae-Ueng et al. [16] decreases more dramatically than that of other two phage DNAs with smaller capsids adopted in the experiment of Leforestier et al. [19]. This shows that there is a size effect in the transition of viral DNA structure with temperature variation, because a larger virus allows DNA chain to be rearranged more loosely and disorderedly after warmed up. According to this result, we suggest that the controversy about the temperature effect of the viral DNA structure f in previous SAXS experiments [16,17,19] is ascribed to size effect. In fact, this size-dependent collective response in DNA structure is a diagram of DNA intra- and inter-strand interactions as shown in Eqs. (1)–(2) [36].

**FIG. 3.**
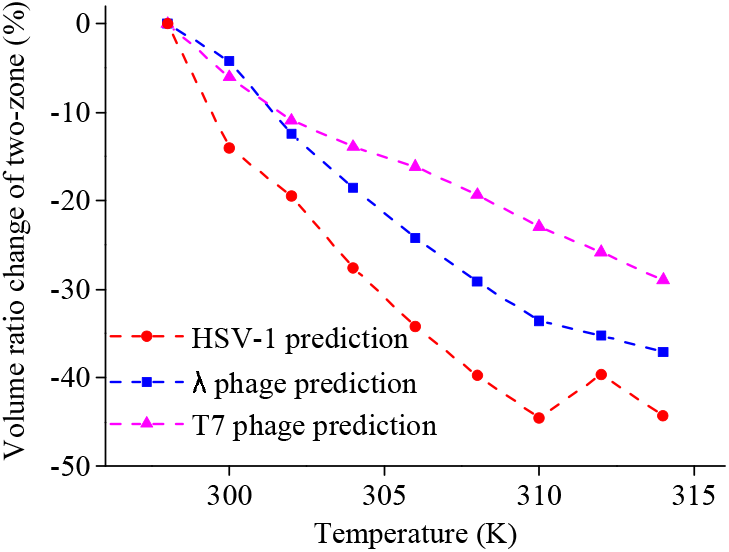
(color online). Two-zone volume ratio change with temperature for three viral DNAs (HSV-1, λ phage and T7 phage) under 0.1 M Na^+^ solution.

Then the bulk stiffness change with the increasing temperature for three viral DNAs can be calculated by Eqs.(1) and (6) with the DNA structural parameters obtained in Figs. 2 and 3. In addition to 0.1 M Na^+^ condition, we also predict the DNA stiffness of HSV-1 under 25mM Mg^2+^ condition [29] to show the salt-dependency of bulk mechanical property. Fig. 4 shows the comparison between the present stiffness predictions for three viral DNAs and the previous experiment of HSV-1 [16]. The prediction of HSV-1 DNA stiffness under 25 mM Mg^2+^ condition is quantitatively consistent with the experimental result, which demonstrates the rationality of the present model. Regardless of virus types and ionic conditions, all the bulk stiffnesses of three viral DNAs decrease monotonously with the increasing temperature. The bulk stiffness of viral DNA in divalent cationic solution is significantly lower than that in monovalent cationic solution, which complies with the general ionic effect of viral mechanical properties [34,38]. The most interesting thing is the dramatic decline of HSV-1 DNA stiffness among the three viruses. This indicates that bulk DNA stiffness also has the size effect, which is another reflection of intermolecular interactions engendering collective responses [36].

**FIG. 4.**
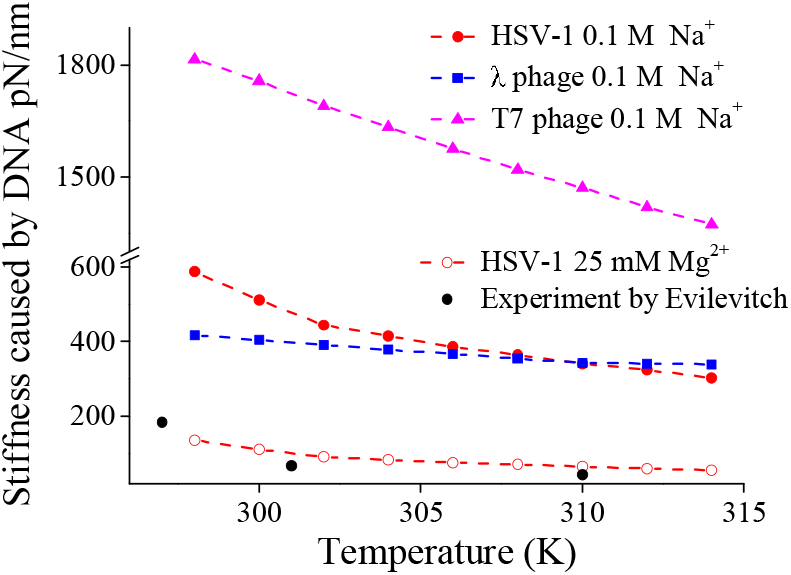
(color online). Bulk stiffness change with temperature for three viral DNAs (HSV-1, λ phage and T7 phage).

Our previous study reveals that a transition of viral DNA structure from order to disorder during DNA ejection can lead to the change of bulk mechanical properties like the ejection force [29]. Therefore, here we suggest that the size effect on the temperature-dependent stiffness of viral DNA in Fig. 4 is presumably related to its structure transition due to temperature variation in Figs. 2 and 3. The results in Figs. 2–4 show that the larger the size of viral capsid, the more drastic change of viral DNA structure during thermal fluctuation; analogous to a sponge, when it expands much larger, its entirety gets softer with more internal voids. In addition, a good matching between the size-dependent change of viral DNA stiffness and that of two-zone volume ratio with temperature for three viruses is redisplayed in Fig. S4 of Supplement 4. This indicates that the size effect of viral DNA stiffness is really related to its structure transition. In the sense of biology, the size effect of stiffness is related to different routes of infection for different viruses. The fluid-like DNA phase under the specific condition is softer than solid-like DNA phase, which increases the mobility of viral genes [17]. Specifically, HSV-1 first adheres to cell surface, then enters cell through endocytosis [39], so the decrease of its stiffness is conducive to viral infection; whereas the bacteriophage is fixed on the bacteria surface and its DNA is ejected into cytoplasm [2], so the slightly changed stiffness implies that there is a sufficient internal pressure in capsid to ensure the ejection.

In conclusion, in order to tackle with the controversy about the temperature-dependency of viral DNA structure in previous experiments [16,17,19], we propose an two-zone structure assumption for viral DNA, and update a mesoscopic energy model of DNA liquid crystalline with two phases and a bulk stiffness model of viral DNA in AFM nano-indentation. These multi-scale models reveal a temperature-dependent correlation between microscopic structure of viral DNA and its bulk stiffness engendered by environment sensitive intermolecular interactions. The present predictions of microscopic structural parameter, mesoscopic free energy and bulk stiffness for viral DNA agree well the relevant experiments. The comparison of three typical viruses (HSV-1, λ phage and T7 phage) shows that thermal rise enlarges interchain spacing, especially in the tighter ordered zone, while the DNA length in the ordered zone decreases. The degree of microstructure transformation between a tighter ordered phase and a looser disordered phase is size dependent, which leads to the size effect of bulk DNA stiffness; the larger the viral size, the more remarkable changes of viral DNA structure and the resultant bulk stiffness. This multiscale temperature-dependent correlation not only clarifies the controversy about viral DNA structure change in recent SAXS experiments [16,17,19], but also elucidates the mechanism of bulk DNA stiffness change in AFM nano-indentation [16,17,38]. The size effect of viral DNA structure and stiffness change enhances the mobility of viral genes, and also reflects different infectious strategies for different sizes of viruses. In the future, this size effect of temperature-dependent microstructure and bulk property deserves to be checked for more viruses and phages by AFM experiments or computer simulations.

This work was supported by Grant Nos. 11772182, 11272193, 10872121 of the National Natural Science Foundation of China and Grant No. 2019-01-07-00-09-E00018 of the Program of Shanghai Municipal Education Commission.

## Supporting information

Supplemental Materials (including Supplement 1, 2, 3, 4)

