## Supplemental Materials (including Supplement 1, 2, 3, 4) for "Size Effect on Structure and Stiffness of Viral DNA during Temperature Variation"

1. The structural parameter change with temperature for two phage DNAs (λ and T7)

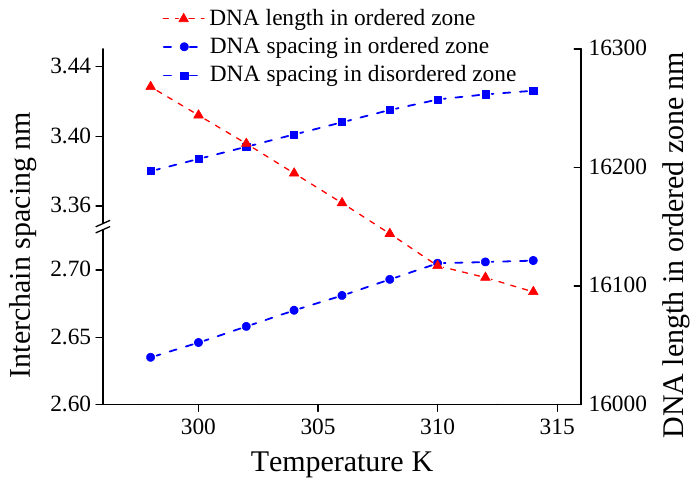

(a)

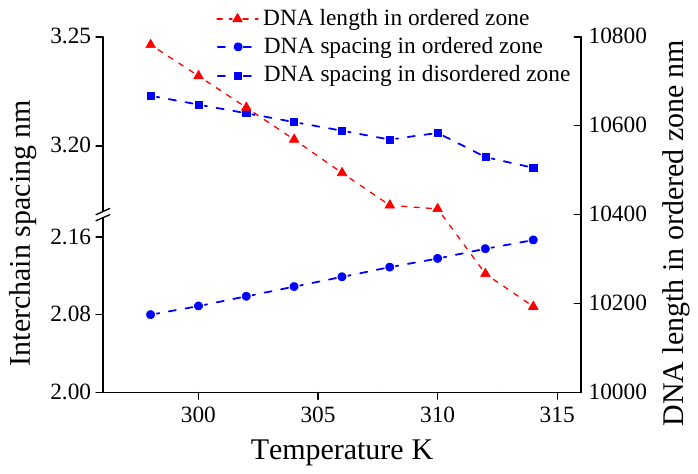

(b)

Fig. S1(b) (color online). Structural parameter change with temperature for two phage DNAs under 0.1 M Na^+^ solution (a) λ phage, (b) T7 phage

2. The effective volume of viral DNA

Fig. S2 shows the effective volume of viral DNA including the hexagonal area around DNA cylinder. The effective volume is written as

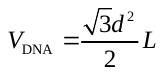
, (S1)

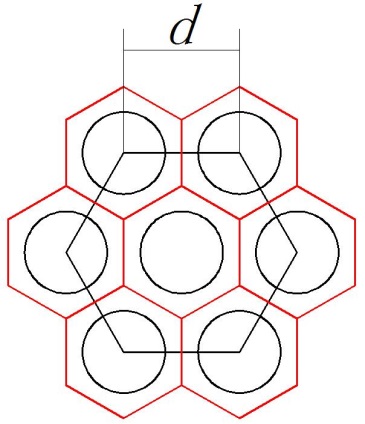

DNA chain

Fig. S2 (color online). Effective volume of viral DNA

3. The bulk stiffness contributed by viral DNA

The bulk stiffness contributed by viral DNA in the process of AFM nano-indentation will be formulated. Assume that there is no energy dissipation during the indentation process, given the force applied to the probe *F* and the indentation depth *D*, all the work done by the probe is converted to the energy change of viral particle, expressed as

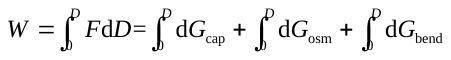
, (S2)

where
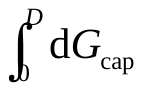
is the energy change contributed by the capsid deformation,
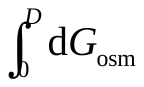
is the energy change contributed by DNA osmotic pressure in capsid,
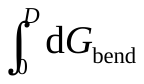
is the energy change contributed by DNA bending elasticity. Taking the derivative of both sides of Eq. (S2) with respect to the indentation depth *D*, the relationship between the compression and the resistance contributed by the virus is obtained as

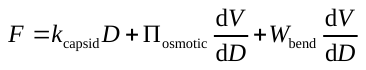
, (S3)

Considering the geometrical changes of the capsid during indentation process [33], the viral particle stiffness is models as

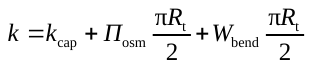
. (S4)

The elastic bending energy density of a DNA chain is written as

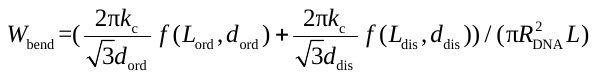
. (S5)

in which *R*_DNA_ is the radius of a DNA cylinder and *R*_DNA_ =1 nm [20].

4. The relative changes of bulk stiffness and two-zone volume ratio with temperature for three viral DNAs

As shown in Fig. S4, all the relative changes of bulk stiffness and two-zone volume ratio for three viral DNAs (HSV-1, λ phage and T7 phage) decrease with the increasing temperature. The changes of bulk stiffness and microstructure of HSV-1 DNA with the largest capsid size are the most drastic, and that of the other two phages with smaller capsid sizes are relative moderately, which shows a remarkable size-dependence. However, there are some of the differences in the correlation between the two changes, for example, the change in the DNA structure of λ phage is greater than that of T7 phage, whereas the change in the bulk DNA stiffness of λ phage is smaller than that of T7 phage. This partial difference indicates that the correlation between microstructure and bulk stiffness of viral DNA is not a simple linear correspondence, and the size effect is not only dependent on the capsid size, but also the packaged DNA length. This also demonstrates the diversity of biological soft matter.

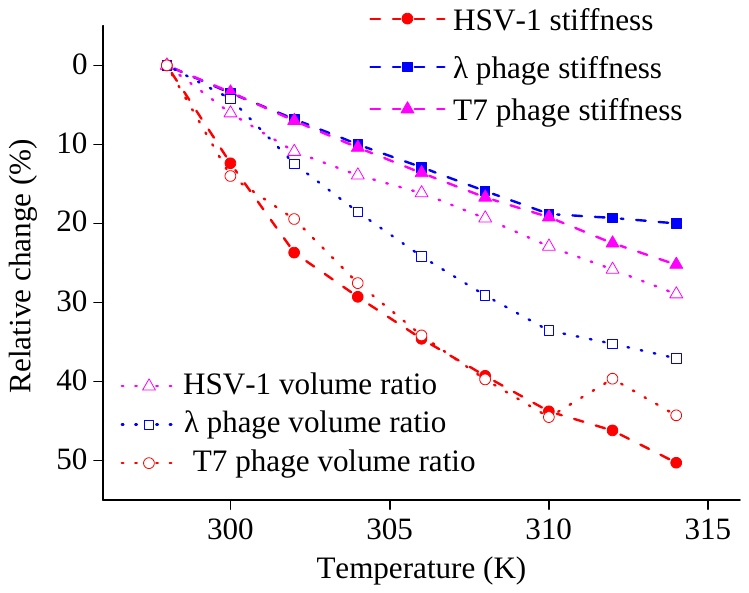

Fig. S4 (color online). Relative changes of bulk stiffness and two-zone volume ratio with temperature for three viral DNAs (HSV-1, λ phage and T7 phage).
